# A model explaining mRNA level fluctuations based on activity demands and RNA age

**DOI:** 10.1101/2020.04.30.069674

**Authors:** Zhongneng Xu, Shuichi Asakawa

## Abstract

Cellular RNA levels are usually fluctuated and are affected by many factors. However, knowledge on the fundamental relationships between RNA abundance, environmental stimuli, and RNA activities is lacking, and the effects of RNA age on RNA levels remain unknown. A model based on activity demands and RNA age was developed to explore the mechanisms of RNA level fluctuations. With single cell time-series gene expression experimental data, we could assess transcription rates, RNA degradation rates, RNA life spans, RNA demands, accumulated transcription amounts, and accumulated RNA degradation amounts by this model. The model could also predict RNA levels under simulation backgrounds, such as stimuli leading to regular oscillations in RNA abundance, stable RNA levels over time being resulted from the long shortage of total RNA activity or uncontrollable transcription, and relationship between RNA levels and protein levels. Given explaining partly mechanisms of some RNA level fluctuations by the present model, further proves and suitable applications are expected.

## Introduction

The mechanisms underlying the establishment of specific RNA levels in a cell are a target of transcriptomic study (Patel et al., 2014; Treutlein et al., 2016; Villani et al., 2017). Detected cellular RNA levels are usually fluctuated (Deng et al., 2014; Briggs et al., 2018; Rodriguez et al., 2019). Studies have provided diverse explanations to the mechanisms of RNA level fluctuations (Elowitz and Leibler, 2000; Chubb et al., 2010; Baldazzi et al., 2012; Bar-Joseph et al., 2012; Stavreva et al., 2012; Briggs et al., 2018; Yamada and Akimitsu, 2019). However, limited transcriptomic techniques meant that some potential fundamental effects on RNA level fluctuations remained unknown.

Little is known about the relationship between supply and demand of RNA. RNA excesses or deficiencies can be harmful (Carper, 1982; Cole et al., 1992; Humpherys et al., 2002; Varambally et al., 2008). Regulation of appropriate RNA levels to meet cellular demands of RNA appear to be necessary to maintain cellular health. Experimental evidence showed that some transcription initiation events result from deterministic factors such as through intrinsic and/or extrinsic signals (Rusak et al., 1990; Stavreva et al., 2009; Hao and O’Shea 2011; Battich et al., 2015). The challenge is to link RNA level fluctuations with the effects of different environmental stimuli, the demands of RNA, and transcription.

RNAs are queued along the DNA template for their production by RNA polymerase. The term “RNA age”, which specify how long the RNAs have existed since their initial transcription, was previously used in our research of a gene expression model (Xu and Asakawa, 2019a) and later Rodriques et al used it in an RNA timestamp analysis (Rodriques et al., 2020). From their birth to their death, RNAs are involved in temporal and spatial biological processes and in modulating various physiological activities (Mata et al., 2005; Moore, 2005; Houseley and Tollervey, 2009). Moreover, the lifespan of RNAs usually range from a few minutes to more than two days (McManus et al., 2015), which are possibly longer than the interval times between two consecutive transcription pulses (pulses usually range from a few minutes to a few hours) (Ozbudak et al., 2002; Chubb et al., 2006; Chubb and Liverpool, 2010; Locke et al., 2011). Thus, transcripts with different RNA ages co-exist in the cellular RNA pool. The relative abundances and expected degradation times of RNAs with different ages are therefore worthwhile considerations in the study of RNA activities and fluctuations of RNA abundance.

Transcription and RNA degradation are key processes that lead to fluctuations in RNA levels (McManus et al., 2015; Yamada and Akimitsu, 2019). The detected cellular RNA levels within an experiment are not RNA amounts transcribed in the experimental period, but RNA abundance, which is RNA accumulation plus RNA transcription minus RNA degradation (Xu and Asakawa, 2019b). Special methods to examine transcription rate and RNA degradation rate have been reported (Tani, et al, 2012; Schwalb, et al, 2016), but they are hard to be used in the normal RNA experiments with the living organisms. Thus, the generally applicable method on quantitative estimation of transcription rate and RNA degradation rate is still lacking.

In the present research, we developed a model based on the demands of RNA, RNA age (which determine the survival time and biological activity of an RNA), transcription, and RNA degradation to explain the mechanism underlying RNA level fluctuations in a cell. We also explored the applicability of the model for analysing dynamic processes between interacting biomolecules, such as the relationship between RNA and protein level fluctuations. The aim is to provide explanations to some of the transcriptomic phenomena and to provide a new perspective for future studies.

## Materials and methods

### Description of the model

A model was built based on the production of biomolecules (including RNAs and proteins) as triggered by the demand of biomolecule activities (DA) and their age-dependant degradation kinetics (Figure 1a). The age of a biomolecule was defined as the length of time the biomolecule existed since its production. The production of biomolecules was induced by stimuli. There was a maximum number of biomolecules at age 0 due to the limits imposed by molecular effects and micro spaces (Stavreva et al., 2012; Battich et al., 2015; Liu et al., 2016). After this time, the biomolecules became mature and aged and were gradually degraded. Percentage of the survival rate was used to describe the degradation of biomolecules. Biomolecule ages were considered to affect their survival and activities of biomolcules. We supposed that DA is a physiological reaction to a stimulus derived from the intrinsic or extrinsic environment and that the DA value reflected the stimulus level. In the model, if the total existing biomolecule activity (TA) was less than the DA, then the production of biomolecules was triggered; otherwise, the production of biomolecules ceased. The biomolecule abundance and biomolecule activity in this model were calculated as follows:

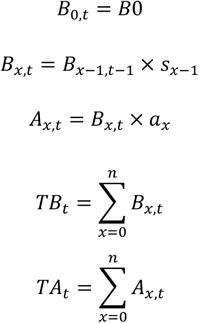

where *B* is the abundance of the biomolecule; *t* is the time; *B*0 is the abundance of biomolecules at age 0; *x* is the age of the biomolecules; *s* is the degradation coefficient (survival percentage in relation to biomolecule abundance); *A* is the activity of the biomolecules; *a* is the activity coefficient; *TB* is the total amount of biomolecules of all ages; *n* is the maximum age of the biomolecules; and *TA* is the total activity of the biomolecules of all ages.

**Figure 1.**
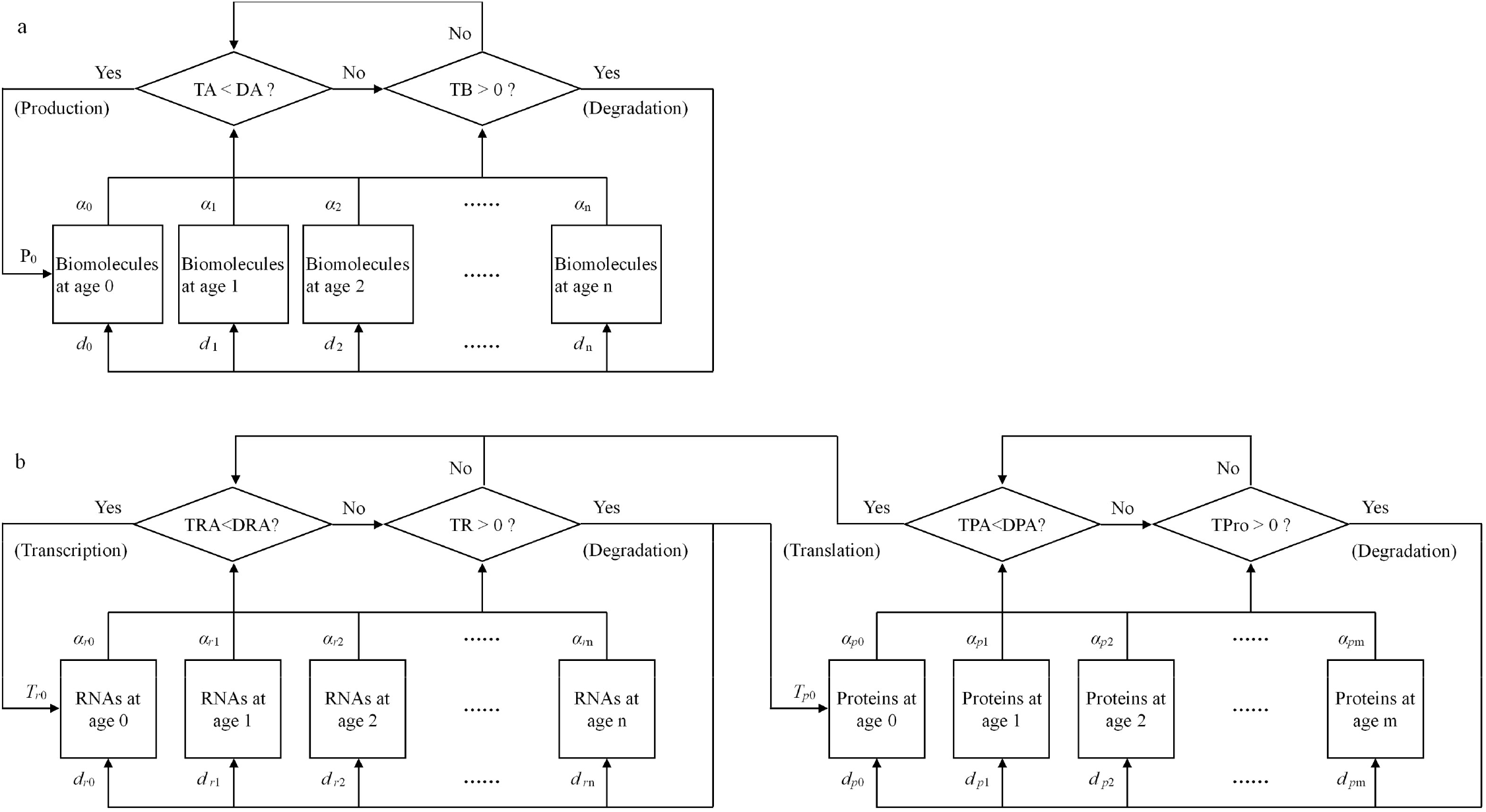
The model of biomolecular level fluctuations. a. A one-unit model of biomolecular dynamics in a cell. The TB represented the total abundance of the biomolecules at all biomolecular ages. The TA represented the total biomolecular activity at all biomolecular ages. The DA represented the demands of biomolecular activity in the cell. The DA was affected by stimuli from intra- and extra-cellular environments, and the difference between the TA and DA determined the trigger or cessation of biomolecule production. *d*_0_, *d*_1_, *d*_2_, …, *d*_n_ were degradation coefficients. *α*_0_, *α*_1_, *α*_2_, …, *α*_n_ were activity coefficients. P0 represented the amount of nascent biomolecule at age 0. b. A two-unit model of the dynamic of RNAs and proteins from a gene in a cell. TR is the total amount of RNA of all ages in the cell. TAR is the total RNA activity of all ages in the cell. DAR is the demand of RNA activity in the cell. ** TPro is the total amount of protein of all ages in the cell. TAP is the total protein activity of all ages in the cell. DAP is the demand of protein activity in the cell. DAP corresponds to stimuli from outside and inside. The difference between TAP and DAP and that between TAR and DAR result in the intensity of producing or/and degradation of proteins and RNAs, respectively. *d*_*r*0_, *d*_*r*1_, *d*_*r*2_, …, *d*_*r*n_ are degradation coefficients of RNA. *d*_*p*0_, *d*_*p*1_, *d*_*r*2_, …, *d*_*p*n_ are degradation coefficients of protein. *α*_*r*0_, *α*_*r*1_, *α*_*r*2_, …, *α*_*r*n_ are activity coefficients of RNA. *α*_*p*0_, *α*_*p*1_, *α*_*p*2_, …, *α*_*p*n_ are activity coefficients of protein. *T*_*r*0_ is the amount of nascent RNA per time. *T*_*p*0_ is the amount of nascent protein per time.

### Simulating experimental data and calculating the parameters

We used the model to simulate experimental data of mRNA level fluctuations in single cells (Table S1) reported in the references (Nuñez et al., 1998; Hao and O’Shea, 2011), calculating the parameters such as transcription rates, RNA degradation rates, RNA demands, RNA life spans, RNA survival rated based on RNA ages, accumulated transcription amounts, and accumulated RNA degradation amounts. The exhaustive search method was used in the simulation and minimum square was the selection standard. Due to reducing the calculation amount of the exhaustive search method, the activity coefficients were set to be 1, thus the RNA activity equalled to the RNA level and the RNA activity demand equalled to the RNA demand. Five RNA ages, which averagely divided an RNA span into four periods, were set. The RNA lifespan, RNA demands, RNA survival rates of each RNA age, transcription amount at the time RNA abundances lower than RNA demands, RNA age distribution of RNA abundance at the beginning of the experiment were set as unknown variables to be assessed. To make the calculation with the exhaustive search method possible, two scenarios were set as followed: if the RNA levels were stable or had periodic fluctuations, we set the stable RNA demand through time; if the RNA levels randomly fluctuated, we used every 4 experimental expression values to assess the variables mentioned and only kept the RNA demands values, then at last we let the RNA demands values as the known variables and assessed other variables. In the exhaustive search method, the lower limit of lifespan of RNA in the yeast cell was set 10 min and the lower limit of lifespan of RNA in the mammalian cell was set 1 h according to the reference (McManus et al., 2015).

### mRNA level fluctuations in a cell under different simulation environments

A one-unit model was used to explain RNA fluctuations in a cell. The values of the following parameters were set to simulate the transcriptional features of a gene: the demand of RNA activity (DRA), the RNA level at age 0, RNA survival rates, and the RNA activity coefficients (Table S2). Intra- and extra-cellular stimuli were incorporated into the DRA. Stable and fluctuating DRA values were both incorporated into the design. The RNA lifespan was standardized and divided into 11 ages, from age 0 to age 10 when all RNAs are completely degraded. The RNA ages influence RNA survival and activity. The four types of relationships between RNA age and RNA survival or RNA activity are: Type A, the parameter values for RNA survival or RNA activity being stable at all RNA ages; Type B, the parameter values descend with increasing RNA age; and Type C, the trend of the parameter values is parabolic; Type D, the parameter values increase with increasing RNA age. To enable comparison of our simulation results with data from elsewhere, the RNA levels and RNA activities were standardized as unitless values. Total RNA levels and total RNA activity levels (TRA) were the outputs of the model simulation.

General patterns of cellular RNA fluctuations may be found in a number of single-cell gene expression studies (Stavreva et al., 2009; Hao and O’Shea, 2011; Deng et al., 2014; Briggs et al., 2018; Rodriguez et al., 2019). We used the data provided by these studies to verify the outputs of the model simulation.

### The relationship between mRNA and protein level fluctuations under different simulation environments

A derivative two-unit model was built to explain the relationship between RNA and protein dynamics (Figure 1b). Extra- and intra-cellular stimuli were incorporated into the demand of protein activity (DPA). If the total protein activity (TPA) formed by the total amount of protein (TPro) was less than the DPA, then the difference between the values was set as a stimulus to produce DRA, thereby driving RNA fluctuations. A set of parameter values in this two-unit model (Table S3) was used to evaluate the dynamic relationship between transcription and translation. The total amount of RNA (TR) and TPro were the outputs of the model simulation.

The relationship between cellular protein levels and mRNA abundance are summarized in the reported data (Liu et al., 2016). We used the data provided by these studies to verify the outputs of the model simulation.

### Programming

The simulation processes were translated into C++ programs (provide if requested), and the simulation outputs (provide if requested) were recorded. Plotting functions in R program were used to draw the curves of the simulation outputs.

## Results

### Explaining mRNA level fluctuations in the experiment data by this model

In the results of simulation on three gene time-series expression (Figure 2), R^2^ of mRNA level of each gene is over 0.89. Transcription rates, RNA degradation rates, RNA demands, RNA life spans, accumulated transcription amounts, and accumulated RNA degradation amounts were calculated. In the results of analysing mRNA level fluctuations of *Saccharomyces cerevisiae* gene HSP26 (Hao and O’Shea, 2011) by the model, life span of RNA of gene HSP26 was 20 minutes, the transcription level at each time was 1.4 units, the rate of survival at RNA age 1 was 0.9, the rate of RNA age 2 was 0.9, and the rate of RNA age 3 was 0.5. In the results of analysing mRNA level fluctuations of *S. cerevisiae* gene YNR014W (Hao and O’Shea, 2011), life span of RNA of gene YNR014W was 20 minutes, the transcription level at each time was 3 units, the rate of survival at RNA age 1 was 0.2, the rate of RNA age 2 was 0.2, and the rate of RNA age 3 was 0. In the results of analysing mRNA level fluctuations of mouse GT1-1 cell gene GnRH (Nuñez et al, 1998), life span of RNA of gene GnRH was 4 hours, the transcription level at each time was 54 units, the rate of survival at RNA age 1 was 0.9, the rate of RNA age 2 was 0.9, and the rate of RNA age 3 was 0.8.

**Figure 2.**
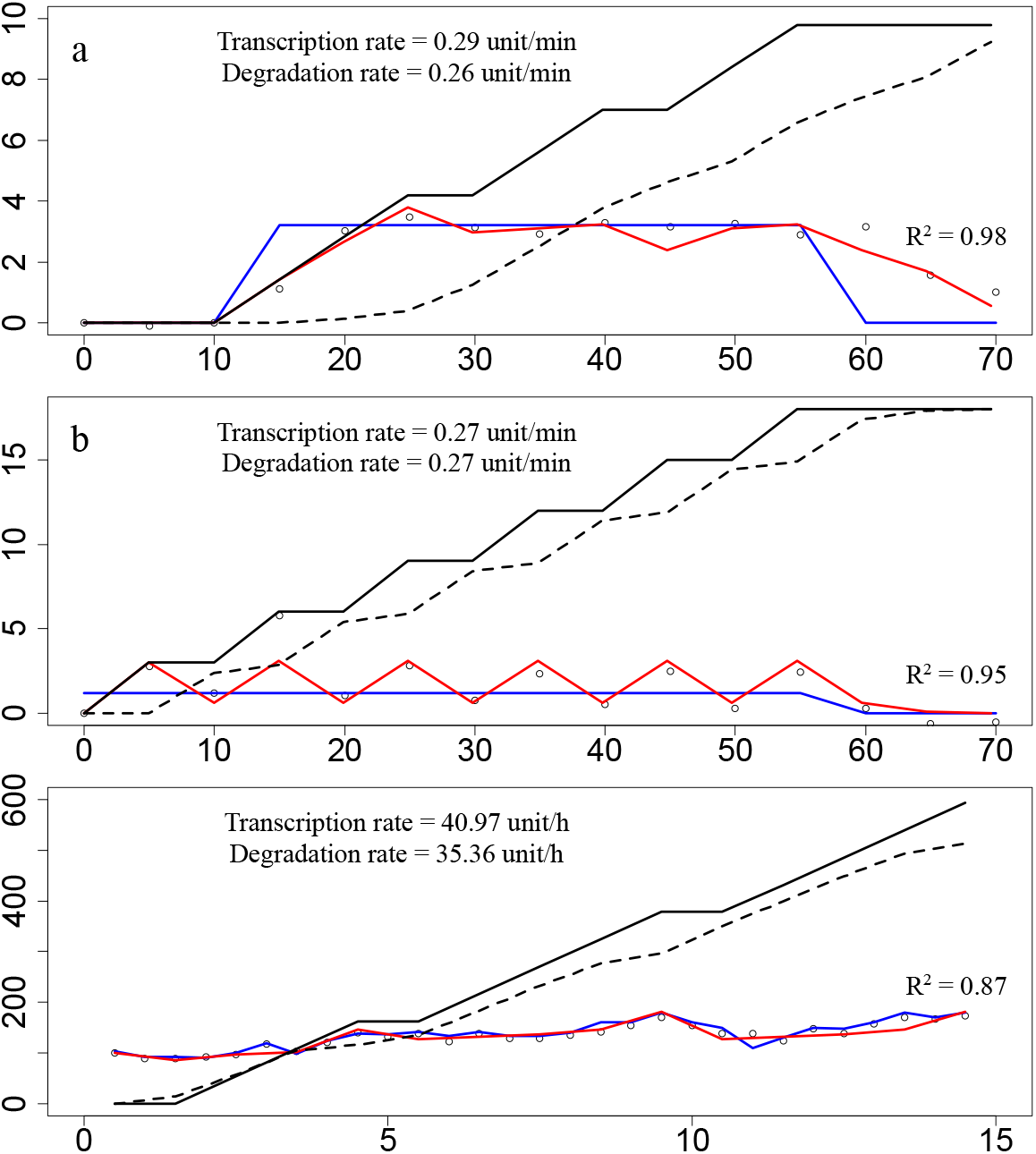
Analysing experiment data of single-cell RNA abundance by the present model. The hollow circles represented the RNA levels of the experiment data from the references (Nuñez et al., 1998; Hao and O’Shea, 2011). The blue line represented the demand on RNA activity (DRA). The red curve represented the RNA level of simulation result of the model. The black solid line represented the accumulative transcription RNA level calculated by the model. The black dashed line represented the accumulative degradation RNA level calculated by the model. a. Analysing mRNA level fluctuations of *Saccharomyces cerevisiae* gene HSP26 by the model. The experiment data were reported by Hao and O’Shea (2011). The X-axis was time, and the unit of time was minute. The Y-axis was RNA level, and the unit was normalized fold change of mRNA level with baseline subtracted. Life span of RNA of gene HSP26 was 20 minutes, the transcription level at each time was 1.4 units, the rate of survival at RNA age 1 was 0.9, the rate of RNA age 2 was 0.9, and the rate of RNA age 3 was 0.5. b. Analysing mRNA level fluctuations of S. *cerevisiae* gene YNR014W by the model. The experiment data were reported by Hao and O’Shea (2011). The X-axis was time, and the unit of time was minute. The Y-axis was RNA level, and the unit was normalized fold change of mRNA level with baseline subtracted. Life span of RNA of gene YNR014W was 20 minutes, the transcription level at each time was 3 units, the rate of survival at RNA age 1 was 0.2, the rate of RNA age 2 was 0.2, and the rate of RNA age 3 was 0. c. Analysing mRNA level fluctuations of mouse GT1-1 cell gene GnRH by the model. The experiment data were reported by Nuñez et al. (1998). The X-axis was time, and the unit of time was hour. The Y-axis was RNA level, and the unit of mRNA level was normalized photonic emissions. Life span of RNA of gene GnRH was 4 hours, the transcription level at each time was 54 units, the rate of survival at RNA age 1 was 0.9, the rate of RNA age 2 was 0.9, and the rate of RNA age 3 was 0.8.

### mRNA level fluctuations in a cell under different simulation environments

Results of the simulation showed dynamic cycles of the total amount of RNA (TR) and the total RNA activity level (TRA) under the condition of stable demands of RNA activity (DRA) (Figure 3), observations that are consistent with the single-cell gene expression pattern reported in genetic laboratory studies (Figure 3a). If TRA was less than DRA, then nascent transcripts were produced. TRA increased even though RNAs were simultaneously degraded. When TRA was higher than DRA, transcription stopped while RNA degradation continued, and TRA decreased until it was again less than the DRA. Age differences in RNA activity and in RNA survival and the RNA level at age 0 affected the cycle lengths of TR and TRA oscillations. Parabolic changes in the RNA activity coefficients caused TRA to respond slowly to DRA compared to the highest RNA activity coefficient of 1, leading to an extension of the cycle length from 5 to 8 RNA ages (Figure 3b and 3c). Introduction of a maturation period for RNA activity, which was incorporated into the parameter design by assigning RNAs lower activities at their early ages (Type C and Type D RNA activity coefficients in Table S1), caused a delay in the TRA peak. Thus, TRA peaks occurred later than TR peaks in the cycling rhythms (Figure 3c). Different DRA values, which were caused by different levels of environmental stimuli, could alter TR oscillation amplitudes and cycling lengths. The wavelength of TR with a DRA of 80 was 7 RNA ages and that with a DRA of 50 was 5 RNA ages. The oscillation amplitude of TR with a DRA of 80 was 96 and that with a DRA of 50 was 85. The difference in TR values with DRA values of 80 and 50 fluctuated dramatically, and TR with a DRA of 80 could be either higher or lower than that with a DRA of 50 (Figure 3d). Using different sampling time might lead to incorrect results (Figure 3e). In the simulation results, the sampling interval of 1 RNA age could obtain the correct oscillatory details, the wave length of 8 RNA ages and the peak of 313 units; the sampling interval of 8 RNA ages led to the stable RNA level of 53 units; the sampling interval of 9 RNA ages led to the wave length of 71 RNA ages; and the sampling interval of 10 RNA ages led to the wave length of 39 RNA ages.

**Figure 3.**
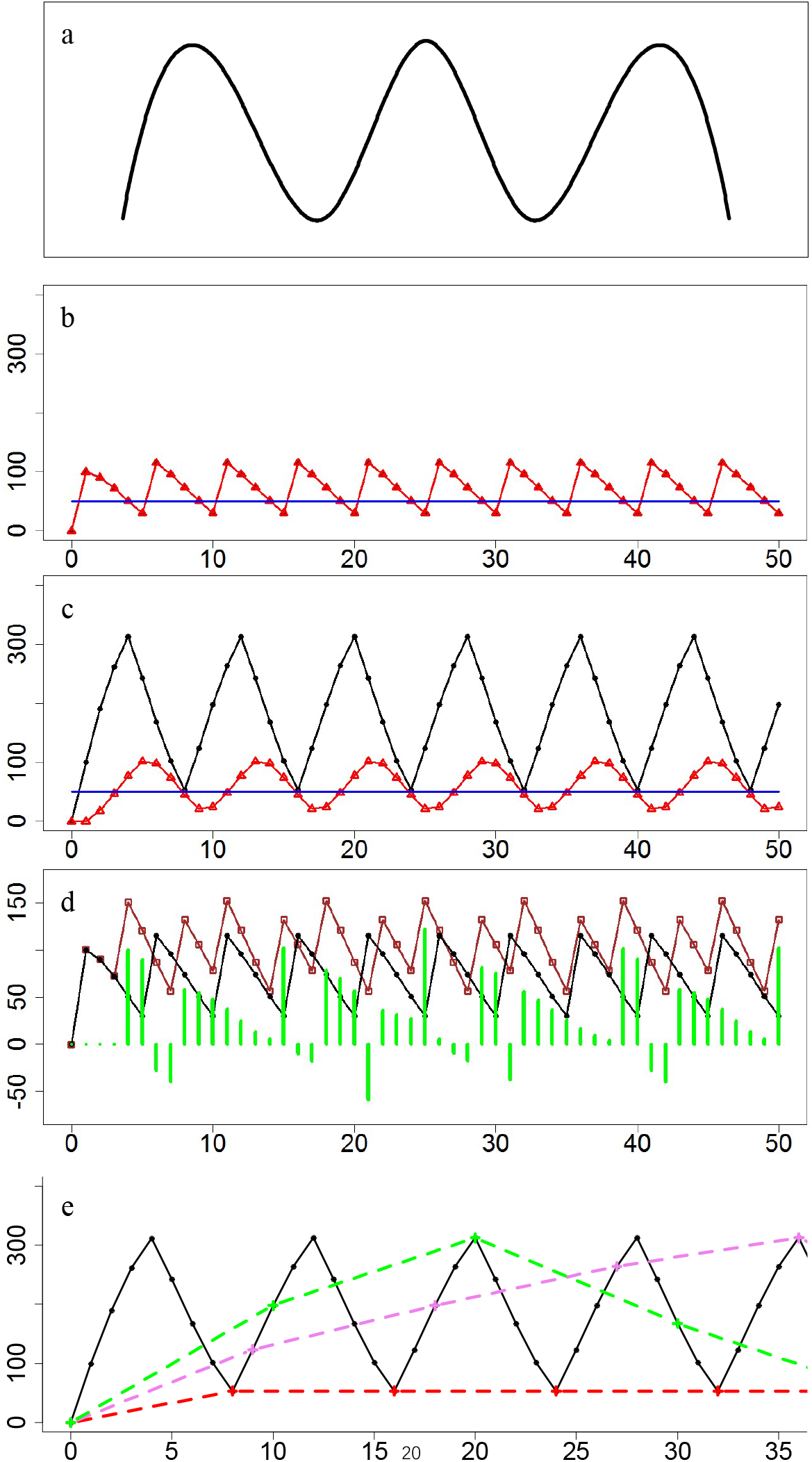
Cycles in RNA abundance dynamics. The blue line represented the demand on RNA activity (DRA), which was here set to 50. The black curve with dots represented the total RNA amount (TR) with a DRA of 50, and the brown curve with hollow squares represented the TR with a DRA of 80. The red curve with triangles represented the total RNA activity (TRA). The green vertical segments represented the differences between the TR with a DRA of 50 and the TR with a DRA of 80. a, Validation data: the oscillatory RNA level reported in the reference (Bar-Joseph et al., 2012). b, The oscillatory RNA with a cycle length of 5 RNA ages. The TR and TRA curves overlapped. (The DRA was set to 50. If the TRA<DRA, then the RNA level at age 0=100; otherwise, the RNA level at age 0=0. The survival rate decreased with older RNA ages, i.e., the Type B RNA survival rate in Table S2. The RNA activity coefficient of each RNA age was 1, i.e., the Type A RNA activity coefficient in Table S2.) c, The oscillations in RNA with shorter cycle length compared to Figure 3b, resulted from lower RNA survival rates. The TR and TRA curves overlapped. (The DRA was set to 50. If the TRA<DRA, then the RNA level at age 0=100; otherwise, the RNA level at age 0=0. The survival rate had a parabolic trend, i.e., the Type C RNA survival rate in Table S2. The RNA activity coefficient of each RNA age was 1, i.e., the Type A RNA activity coefficient in Table S2.) d, The differences between the TR with a DRA of 50 and the TR with a DRA of 80. The green vertical segments were given by: the TR with a DAR of 80 minus the TR with a DAR of 50. (If the TRA<DRA, then the RNA level at age 0=100; otherwise, the RNA level at age 0=0. The survival rate decreased with older RNA ages, i.e., the Type B RNA survival rate in Table S2. The RNA activity coefficient of each RNA age was 1, i.e., the Type A RNA activity coefficient in Table S2.) e, Losing oscillatory details in the results by using different sampling time. The black solid line means the sampling interval of 1 RNA age. The red dash line means the sampling interval of 8 RNA ages. The violet dash line means the sampling interval of 9 RNA ages. The green dash line means the sampling interval of 10 RNA ages. (The DRA was set to 50. If the TRA<DRA, then the RNA level at age 0=100; otherwise, the RNA level at age 0=0. The survival rate decreased with older RNA ages, i.e., the Type B RNA survival rate in Table S2. The RNA activity coefficient of each RNA age was 1, i.e., the Type A RNA activity coefficient in Table S2.)

Environmental variations resulted in changes in the DRA. Cyclic DRAs could lead to new cycles of RNA dynamics (Figure 4). When the DRA was changed from a stable value of 50 (Figure 3a) to a cyclic fluctuation between 50 for 5 RNA ages and 150 for 5 RNA ages, the cycle length of TR fluctuation could change from 5 RNA ages to 30 RNA ages (Figure 4b). The cycle lengths of TR and/or TRA were sometimes the same as those of cyclic DRA (Figure 4b and 4c), an outcome that matched the single-cell gene expression pattern reported in other studies (Figure 4a). An aperiodic DRA could eliminate the cyclic rhythms observed in RNA dynamics (Figure 4d). If stable periods of cyclic DRA were sufficiently long, then the sub-cycles of RNA dynamics may be expected to occur during these periods. When DRA alternated between 50 for 25 RNA ages and 150 for 25 RNA ages, three cycles in two hierarchies were observed. The sub-cycle of 5 RNA ages occurred in stable periods with a DRA of 50, that of 7 RNA ages occurred in stable periods with a DRA of 150, and there was a global cycle of 50 RNA ages over time (Figure 4c). With fluctuating DRAs, several outcomes were possible; for example, TRA could correspond to DRA fluctuations (Figure 4b and Figure 4c) or it could correspond to a lower DRA without reaching a higher DRA (Figure 4d). If fluctuations in the DRA maintained regular cycles, then TR and TRA were typically driven into regularly cycling dynamics.

**Figure 4.**
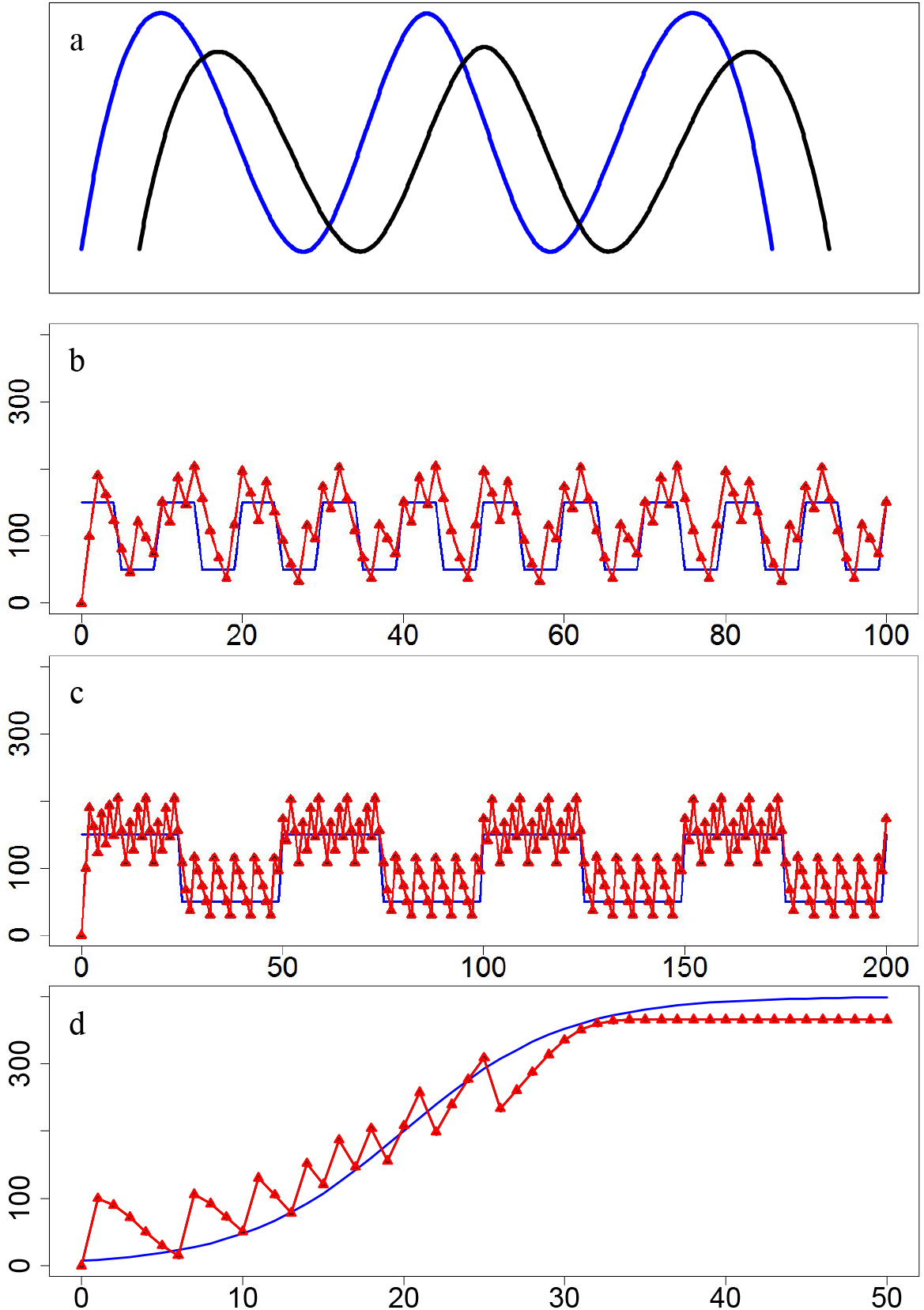
Environmentl variations affecting RNA fluctuations. The blue line represented the demand on RNA activity (DRA). The black curve represented the total amount of RNA (TR). The red curve with triangles represented the total RNA activity (TRA). a, Validation data: the oscillatory RNA level with a oscillatory environmental level in the reference (Stavreva et al., 2012) b, The cycle length of the TR was different from the cycle length of the DRA. The TR curve and the TRA curve overlapped. (The DRA was rotated between 50 for 5 RNA ages and 150 for 5 RNA ages. If the TRA<DRA, then the RNA level at age 0=100; otherwise, the RNA level at age 0=0. The survival rate decreased with older RNA ages, i.e., the Type B RNA survival rate in Table S2. The RNA activity coefficient of each RNA age was 1, i.e., the Type A RNA activity coefficient in Table S2.) c, The cycle lengths of the TR and DRA were the same, and there were local sub-cycles in the global cycles. The TR curve and the TRA curve overlapped. (The DRA was rotated between 50 for 25 RNA ages and 150 for 25 RNA ages. If the TRA<DRA, then the RNA level at age 0=100; otherwise, the RNA level at age 0=0. The survival rate decreased with older RNA ages, i.e., the Type B RNA survival rate in Table S2. The RNA activity coefficient of each RNA age was 1, i.e., the Type A RNA activity coefficient in Table S2.) d, An aperiodic DRA. The TR curve and the TRA curve overlapped. (The DRA was a logistic function: DRA=400/(1+e^4-0.2×time^). If the TRA<DRA, then the RNA level at age 0=100; otherwise, the RNA level at age 0=0. The survival rate decreased with older RNA ages, i.e., the Type B RNA survival rate in Table S2. The RNA activity coefficient of each RNA age was 1, i.e., the Type A RNA activity coefficient in Table S2.)

In some cases, TR and TRA gradually changed to reach stable levels over time (Figure 5), an observation consistent with the reported data (Figure 5a). If transcription continued despite a limiting DRA, i.e. transcription was unregulated, then TR and TRA were adjusted via transcription e and degradation until they reached an equilibrium. (Figure 5b). When TRA was less than DRA over time, TR and TRA values also gradually stabilized (Figure 5c and 5d). On the one hand, if RNA activity coefficients were low, then TRA was also low in spite of a high TR. Under such conditions, despite a low DRA, TRA was less than DRA, and straight lines in TR and TRA subsequently formed (Figure 5c). On the other hand, DRAs that were too high could make it impossible for TRA to reach the DRA (Figure 5d).

**Figure 5.**
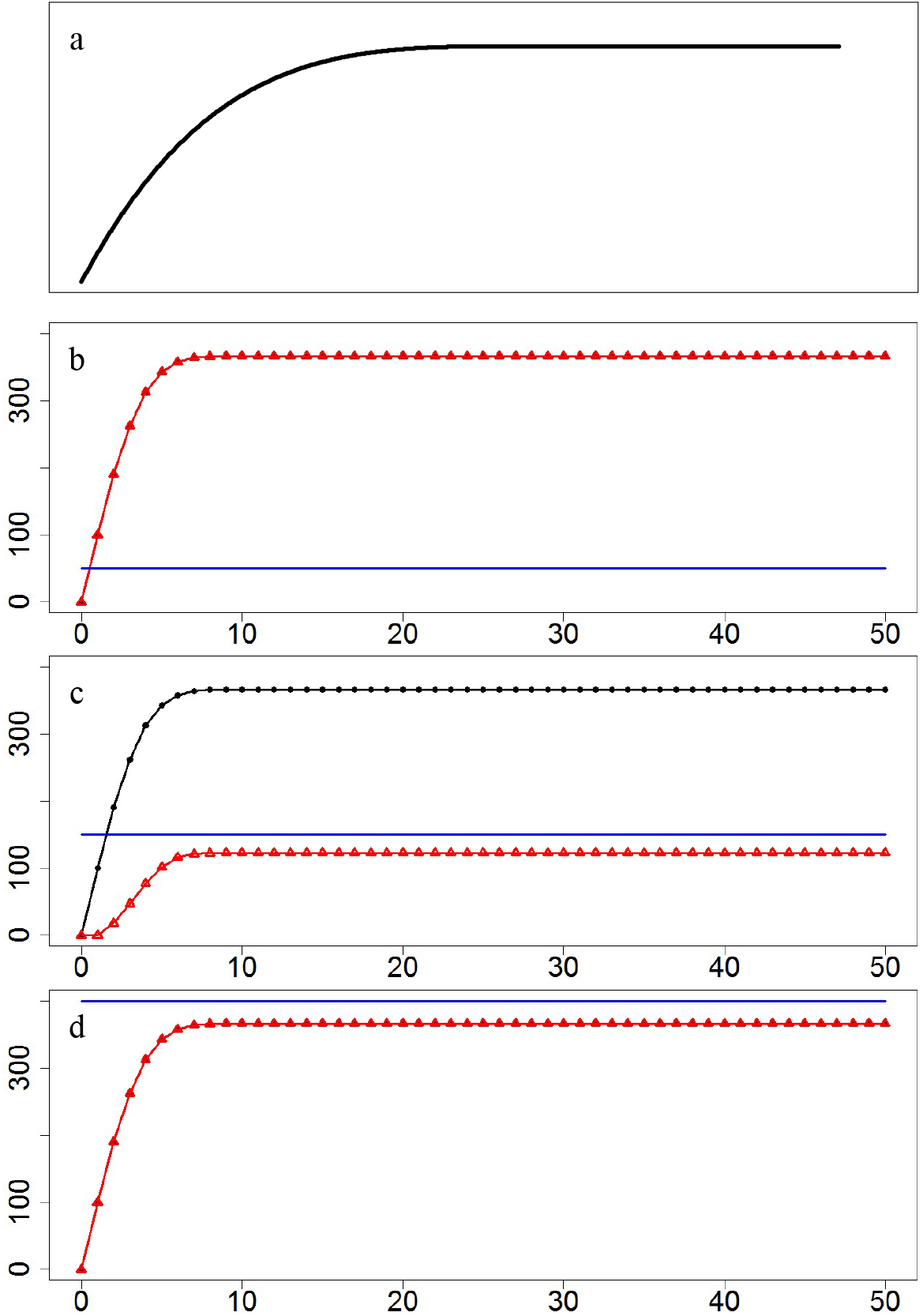
The phenomena of stable RNA levels. The blue line represented the demand on RNA activity (DRA). The black curve represented the total amount of RNA (TR). The red curve with triangles represented the total RNA activity (TRA). a, Validation data: the stable RNA level reported in the references (Bar-Joseph et al., 2012; Hansen and O’Shea, 2015). b, Transcription was unregulated. The TR curve and the TRA curve overlapped. (The DRA was 50. The level of transcripts at age 0 was 100 per time in all periods of the experiment. The survival rate decreased with older RNA ages, i.e., the Type B RNA survival rate in Table S2. The RNA activity coefficient of each RNA age was 1, i.e., the Type A RNA activity coefficient in Table S2.) c, TRA that is too low. (The DAR was 150. If the TRA<DRA, then the RNA level at age 0=100; otherwise, the RNA level at age 0=0. The survival rate decreased with older RNA ages, i.e., the Type B RNA survival rate in Table S2. The RNA activity coefficient had a parabolic trend, i.e., the Type C RNA activity coefficient in Table S2.) d, DRA that is too high. The TR curve and the TRA curve overlapped. (The DRA was 400. If the TRA<DRA, then the RNA level at age 0=100; otherwise, the RNA level at age 0=0. The survival rate decreased with older RNA ages, i.e., the Type B RNA survival rate in Table S2. The RNA activity coefficient of each RNA age was 1, i.e., the Type A RNA activity coefficient in Table S2.)

### Relationship between RNA and protein dynamics

In the two-unit model, the levels of protein and RNA were driven by the demand of protein activity (DPA) (Figure 6). When DPA fluctuated over time, TR and TPro also followed similar trends (Figure 6a). When the DPA value was stable, the total amount of protein (TPro) could undergo fluctuated cycles, accompanied by fluctuated cycles in TR that resulted from DRA cycling (Figure 6b). If the total protein activity (TPA) could not reach the DPA over time, then the TPro changed to a constant value. However, two trends of change may occur. In one case, the TRA could exceed the DRA, resulting in changes in the dynamic cycles of TR (Figure 6c); when TRA was less than DRA over time, then the TR value also stabilizes (Figure 6d).The relationship between RNA and protein dynamics patterns is consistent with that reported in other studies (Figure 6e, Figure 6f, Figure 6g).

**Figure 6.**
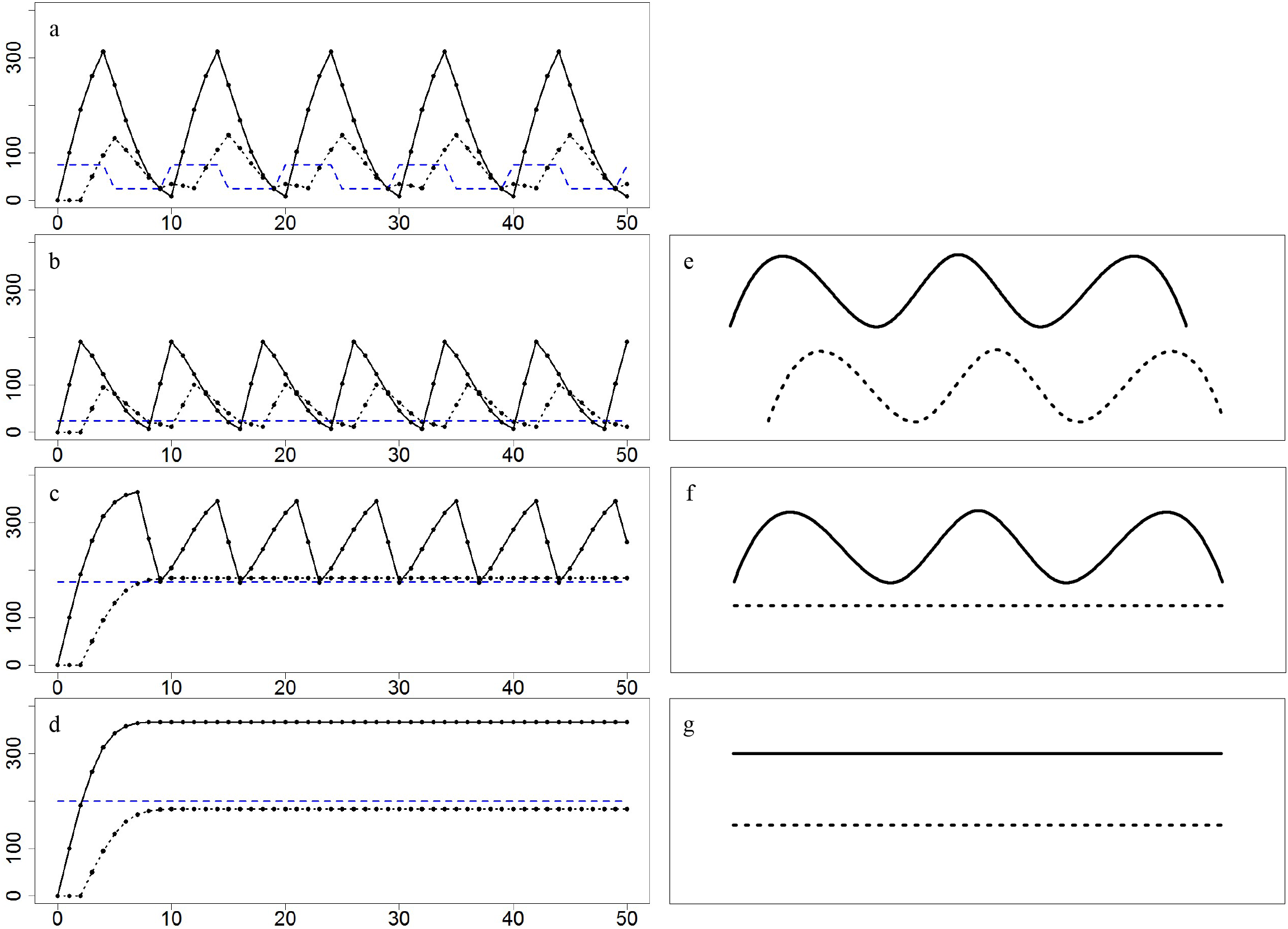
Relationship between RNA and protein dynamics. Four figures in the left side are simulation data: the blue dashed line represented the demand on protein activity (DPA), the black curve with dots represented the total amount of RNA (TR), the black dashed line with dots represented the total amount of protein (TPro), and the values of the parameters were given in Table S3. The three figures in the right side are the valid data: the black curve represented the total amount of RNA (TR), the black dashed line represented the total amount of protein (TPro), and the data were from the previous studies (Liu et al., 2016). a, Cycling TR and cycling TPro under the condition of cycling DPA. (DPA rotated between 50 and 150.) b, Cycling TR and cycling TPro under the condition of stable DPA. (DPA=25) c, Cycling TR and stable TPro under the condition of stable DPA. (DPA=175) d, Stable TR and stable TPro under the condition of stable DPA. (DPA=200) e, Cycling TR and cycling TPro in the reference (Liu et al., 2016) f, Cycling TR and stable TPro in the reference (Liu et al., 2016) g, Stable TR and stable TPro in the reference (Liu et al., 2016)

## Discussion

### Significance of mRNA fluctuations

Dynamic cycling of total biomolecule amount (TB) either occurred under conditions of stable DAs over time (Figure 3c) or were imbedded in the stable phase of the global fluctuations (Figure 4c). If a stable DA originates from a basic cellular requirement, then intrinsic circular rhythms may come into being. Circular changes of environmental stimuli form circular DAs, leading to some TBs with the same rhythms (Figure 6a), a result consistent with reports of RNA rhythms induced by environmental elements such as light, food, hormone signals, and cell-cell communication (Rusak et al., 1990; Stephan 2002; Stavreva et al., 2009; Stavreva et al., 2012). Therefore, the intrinsic circular rhythms of important biomolecules could act as environmental stimuli that affect other biomolecules in establishing biological rhythms in cells, tissues, or even whole organisms. This scenario may also explain some aspects of the mechanism underlying biological rhythms (Hogenesch and Ueda, 2011; Partch et al., 2014).

A positive feedback loop coupled with negative feedback factors partly explains the mechanism of the phenomenon of circadian fluctuations in biomolecules (Elowitz and Leibler, 2000; Sevim et al., 2010). In our model, environmental stimuli such as DRA replaced the positive feedback mechanism to trigger transcription, and transcription stopped when DRA was removed. DRA determined the straight, circadian, or irregular trends in TR. In cases when TRA was less than DRA over time, TR reached a certain value and stabilized over time (Figure 5), and this phenomenon may hold important biological and medical significance. For instance, if a specific RNA is maintained at a stable level, the cell may fail to meet the demands of RNA or its gene expression becomes unregulated. This could result in a specific RNA being selected as a candidate target in the study of gene function, disease diagnosis, drug design, or other endeavours.

### Assess of RNA metabolic parameters and RNA age distribution

RNA metabolic parameters, such as net RNA production amount and net RNA degradation amount, have important biological and medical information, but they are, limited by the technological methods, difficult to be measured in human and living organisms. RNA abundance data have been greatly produced in the past decades, but directly taking RNA abundance as net RNA production amount can easily result in incorrect judgements (Xu and Asakawa, 2019). Given sufficient time series RNA abundance data, the present model successfully estimated transcription rates, RNA degradation rates, RNA demands, RNA life spans, accumulated transcription amounts, accumulated RNA degradation amounts, etc. (Figure 2). In some cases, the estimated values of RNA metabolic parameters calculated by the present model can provide information enough for biological and medical purposes. Some parameters, e.g., the demand of RNA activity (DRA), of the model needed reasonable explanation. In the expression of gene HSP26 (Figure 2a) and gene YNR014W (Figure 2b), DRAs were closely related to the exogenous stimulus signals of 1-NM-PP1 (Hao and O’Shea, 2011). Of course, the detail biological significance of a DRA is sometimes difficult to estimate due to limited information regarding the stimuli, such as the analysis results of mouse GT1-1 cell gene GnRH (Figure 2c).

RNA age affected the RNA degradation rates, which determined whether the TR exhibited cycling as well as what shapes of the response curves of RNA fluctuations. Age-dependent differences in RNA degradation determined the maxima of TR, which in turn affected whether TRA met the DRA. When low RNA survival rates led to rapid TR decreases, the wavelengths of the fluctuation curves were lower (Figure 3c). Although some knowledge already exists regarding the process and effects of RNA ageing, such as RNA post-transcriptional processing and the identification of RNAs transcribed in series transcription (Houseley and Tollervey, 2009; Schwalb et al., 2016), further research on the finer details is needed.

How to measure RNAs ages is a new question. An RNA timestamps approach is used to infer the age of individual RNAs triggered by the same promoter (Rodriques et al., 2020), but this method cannot directly measure the levels of RNA with different ages at the same time. We think that labeled method can solve this problem. RNAs could be labeled, and the labeled RNAs could be isolated and sequenced (Tani et al., 2012; Schwalb et al., 2016). Suppose that several treatment groups were set in the simulation experiment, and the cells of each group were exposed to the labeled media at different time (Figure 7). The levels of RNA at a certain RNA age were calculated from the differences between labeled RNA levels in different groups. The results of the experiment for measuring RNA age structures could be predicted by the present model. In the condition of stable RNA levels, the cell will have the RNAs with consecutive ages, and the level of RNA at each age is in dynamic equilibrium state (Figure 7b). In the condition of fluctuating RNA levels, the cell will have the RNAs with inconsecutive ages, because RNAs are not produced at the time when TRA is over DRA; the level of RNA at each age is fluctuating (Figure 7c). Further related experiments are expected to measure RNA age distribution.

**Figure 7.**
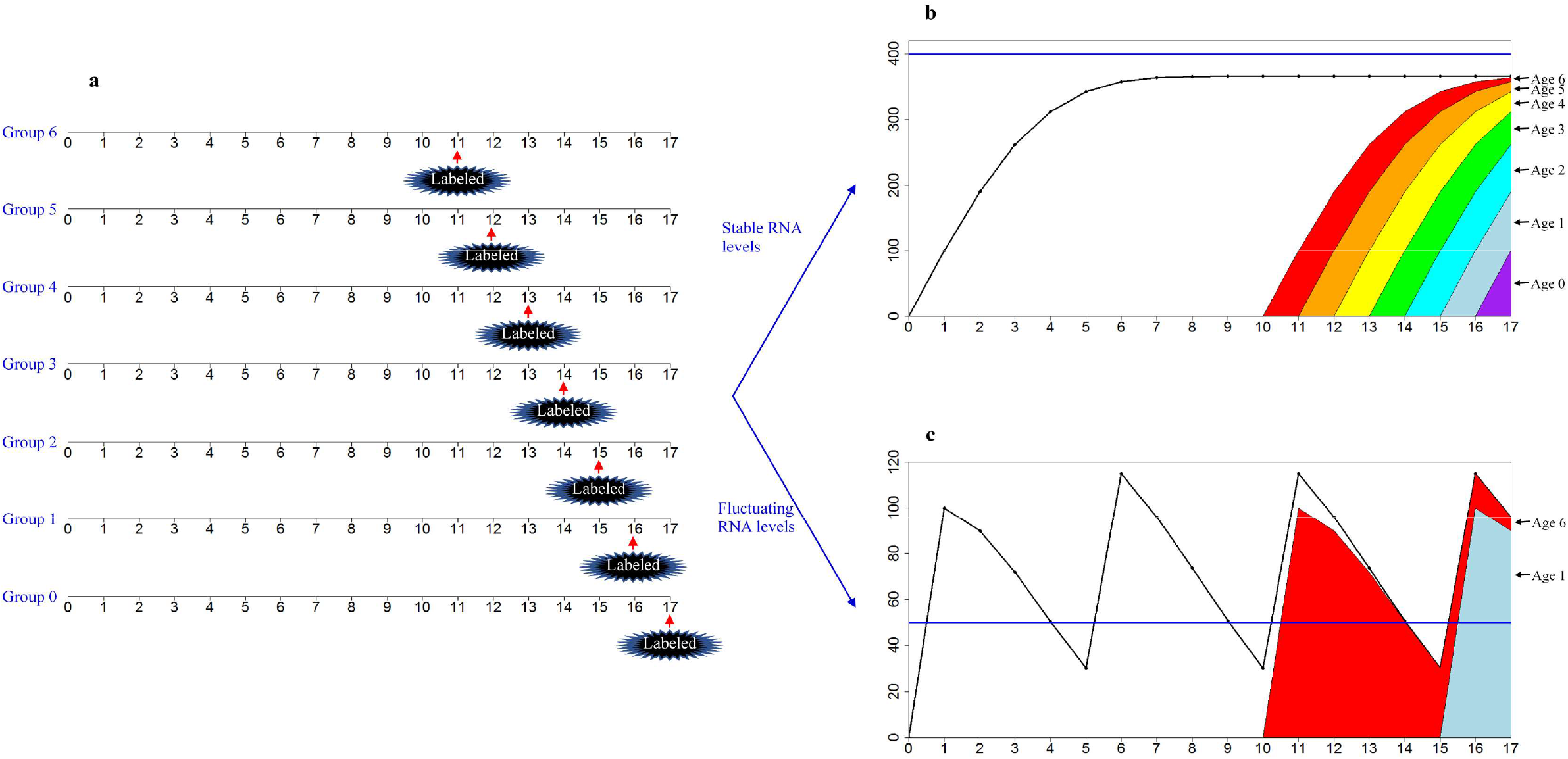
Measurement of RNA age structure. a. Design of the experiment. 6 groups were set in the simulation experiments. Cells were cultured in the non-labeled normal media for some time, and were exposed to labeled media at time 11 in Group 6, at time 12 at Group 5, at time 13 at Group 4, at time 14 at Group 3, at time 15 at Group 2, at time 16 at Group 1, and at time 17 at Group 0. Samples were collected at time 0 to time 17, and RNA levels (of one gene or some genes) were detected. At the time of exposure in labeled media and later times, non-labeled RNAs and labeled RNAs were separated out and detected separately. The RNA levels of a certain RNA age at a certain time were obtained from the differences between labeled RNA levels in two near groups. Take RNA levels at time 17 as an example, RNA levels of Age 6 = labeled RNA levels in Group 6 – labeled RNA levels in Group 5, RNA levels of Age 5 = labeled RNA levels in Group 5 – labeled RNA levels in Group 4, RNA levels of Age 4 = labeled RNA levels in Group 4 – labeled RNA levels in Group 3, RNA levels of Age 3 = labeled RNA levels in Group 3 – labeled RNA levels in Group 2, RNA levels of Age 2 = labeled RNA levels in Group 2 – labeled RNA levels in Group 1, RNA levels of Age 1 = labeled RNA levels in Group 1 – labeled RNA levels in Group 0, and RNA levels of Age 0 was labeled RNA levels in Group 0. b. RNA age structures in the condition of stable RNA levels. The blue line represented the demand on RNA activity (DRA). The black curve with dots represented the total amount of RNA (TR). Colors indicated levels of RNAs labeled at different times, and the RNA levels in the groups whose RNAs labeled at latter times covered those in the groups whose RNAs labeled at earlier times. The RNA level of Age 6, Age 5, Age 4, Age 3, Age 2, Age 1, and Age 0 was 6 (=364-358), 15 (=358-343), 31 (=343-312), 50 (=312-262), 72 (262-190), 90 (190-100), and 100, respectively. (The DRA was 400. If the total RNA activity<DRA, then the RNA level at age 0=100; otherwise, the RNA level at age 0=0. The survival rate decreased with older RNA ages, i.e., the Type B RNA survival rate in Table S2. The RNA activity coefficient of each RNA age was 1, i.e., the Type A RNA activity coefficient in Table S2.) c. RNA age structures in the condition of fluctuating RNA levels. The blue line represented the demand on RNA activity (DRA). The black curve with dots represented the total amount of RNA (TR). Colors indicated levels of RNAs labeled at different times, and the RNA levels in the groups whose RNAs labeled at latter times covered those in the groups whose RNAs labeled at earlier times. The RNA level of Age 6, Age 5, Age 4, Age 3, Age 2, Age 1, and Age 0 was 6 (=96-90), 0 (=90-90), 0 (=90-90), 0 (=90-90), 0 (=90-90), 90 (90-0), and 0, respectively. (The DRA was set to 50. If the total RNA activity<DRA, then the RNA level at age 0=100; otherwise, the RNA level at age 0=0. The survival rate decreased with older RNA ages, i.e., the Type B RNA survival rate in Table S2. The RNA activity coefficient of each RNA age was 1, i.e., the Type A RNA activity coefficient in Table S2.)

### Effects of RNA fluctuations on analysis of transcriptomic data

Fluctuations in cellular RNA abundance can introduce challenges for establishing sampling times and for making comparisons between treatment and control groups. If the RNA levels were stable (Figure 5), sequential sampling, or even one-time sampling, could yield reasonable results. When RNA levels oscillate, the measurement of one-time samples, and even sequential sampling, could yield random results. At the early stages of an experiment, the amount of RNA in cells before the start of the experiment could markedly affect the results (Figure 3, Figure 4, Figure 5); after this point, the levels of the biomolecules were regular. If the levels were oscillatory at the regular stages, then the results would depend on the design of the time intervals for sequential sampling (Figure 3e). When the sampling interval was equal to the wavelength of the cycling curve of the biomolecule, the results of the experiment appeared as a level line on the curve. Perfect and thorough sampling should cover all the feature points of an oscillatory curve. Comparing the treated groups with the controls is the golden rule in biological experiments when the measured values are nearly constant (Figure 5), and statistical comparison between groups is straightforward. In some cases of cyclical biomolecule abundance, however, the levels of biomolecules experiencing high environmental pressures could sometimes be either higher or lower than those experiencing low environmental pressures (Figure 3d). Indeed, in our study, environmental pressures affected the cycle length and vibration amplitudes of biomolecule abundance dynamics (Figure 3d).

It was necessary to distinguish between transcriptional pulses and pulse-like fluctuations in RNA abundance. When a transcriptional pulse occurs, RNA levels increase abruptly. At this point, if rapid degradation follows, RNA abundance drops dramatically and a pulse-like curve in the RNA abundance is formed. Without degradation, a single pulse leads to a horizontal line in the RNA abundance curve, and successive pulses lead to an ascending curve. Thus, transcriptional pulses do not certifiably lead to pulse-like curves for RNA abundance.

### Relationship between RNA and protein fluctuations

If the details of the parameters used in this model were known, then protein levels could (to some extent) be calculated from the RNA levels, and vice versa. Unfortunately, collecting enough information on the many parameters required is not easy, and most of them have not been studied in detail. Protein/mRNA ratios or translation rates of specific genes were previously shown to be constant, and protein levels could be predicted from the mRNA levels (Schwanhäusser et al., 2011; Wilhelm et al., 2014); however, these conclusions had been questioned (Fortelny et al., 2017). In the present research, protein/RNA ratios likely fell, in some cases, within narrow ranges or were even constant, especially under the conditions when the TRA and TPA didn’t meet the requirements (Figure 6d). The cycling levels of RNAs and proteins, however, formed a range of protein/RNA ratios from zero to infinity. Although the protein levels were stable, there were probably varying protein/RNA ratios due to fluctuating RNA levels (Figure 6c). Thus, the application of protein/RNA ratios needed the consideration of the relationship between RNA and protein fluctuations.

## Conclusion

Upon the introduction of biomolecule ages and the demands of biomolecule activity to our model, the cooperation between production and degradation could explain the mechanism underlying cellular RNA fluctuations. The important transcriptomic parameters, such as transcription rates and degradation rates, of time-series gene expression data could be calculated by using this model.

Some new hypotheses were obtained from the analysis results under different simulation environments. Different treatment levels of biomolecule ages and activity could change the wavelength and oscillation amplitude of oscillating RNA or protein levels. Thus, the establishment of sampling times and the result obtained based on typical assumptions regarding the parameters of genetic up- and down-regulation in transcriptomic experiments may be considered carefully. Stable abundance of a specific RNA in a cell probably indicates that the total activities of the RNA could not meet the cellular requirements or that the production of the RNA was unregulated.

This information enriches existing knowledge. Therefore, biological investigations into biomolecule age and biomolecule activity demands and detailed value assignment for specific models require further research.

## Author contributions

Z.X. and S.A. designed the research. Z.X. built the model, conducted the simulation, and wrote the paper. Z.X. and S.A. reviewed the paper.

## Acknowledgements

We thank Dr. Ken Chan for his critical review of our work. X.Z. was supported by the Ocean and Fishery Special Fund Project of Guangdong Province for Technology Extension (grant number 2017A0010).

## Supplementary Tables

**Table S1.** Experiment data of single cell RNA abundance

| Gene name | Experiment time | RNA abundance* | References |
| --- | --- | --- | --- |
| HSP26<br>(from *Saccharomyces cerevisiae*) | (Unit: min)<br>0, 5, 10, 15, 20, 25, 30,<br>35, 40, 45, 50, 55, 60, 65,<br>70 | (Standardized unit)<br>0, -0.1, 0, 1.12, 3.02, 3.47,<br>3.14, 2.91, 3.29, 3.16,<br>3.26, 2.89, 3.16, 1.57,<br>1.01 | Hao and O'Shea, 2011 |
| YNR014W<br>(from *S. cerevisiae*) | (Unit: min)<br>0, 5, 10, 15, 20, 25, 30,<br>35, 40, 45, 50, 55, 60, 65,<br>70 | (Standardized unit)<br>0, 2.78, 1.19, 5.78, 1.07,<br>2.81, 0.77, 2.32, 0.51,<br>2.48, 0.30, 2.42, 0.30,<br>-0.6, -0.54 | Hao and O'Shea, 2011 |
| GnRH<br>(from mouse GT1-1 cell) | (Unit: h)<br>0.5, 1, 1.5, 2, 2.5, 3, 3.5,<br>4, 4.5, 5, 5.5, 6, 6.5, 7,<br>7.5, 8, 8.5, 9, 9.5, 10,<br>10.5, 11, 11.5, 12, 12.5,<br>13, 13.5, 14, 14.5 | (Unit: normalized<br>photonic emissions))<br>100, 89, 88.5, 92.5, 96.5,<br>117.5, 103, 120.5, 140,<br>132.5, 138, 122, 138.5,<br>129, 128.5, 134.5, 141.5,<br>154, 171, 154, 138, 138.5,<br>124.5, 147.5, 138.5, 157,<br>170, 167.5, 173 | Núñez et al., 1998 |

\*RNA abundance values were calculated by drawing grids to analyse the original figures in the papers.

**Table S2.** A set of parameter values used in the model to simulate RNA dynamics

|  | Type | RNA age |  |  |  |  |  |  |  |  |  |  |
| --- | --- | --- | --- | --- | --- | --- | --- | --- | --- | --- | --- | --- |
|  |  | 0 | 1 | 2 | 3 | 4 | 5 | 6 | 7 | 8 | 9 | 10 |
| RNA activity coefficient | A | 1 | 1 | 1 | 1 | 1 | 1 | 1 | 1 | 1 | 1 | 1 |
|  | B | 1 | 0.9 | 0.8 | 0.7 | 0.6 | 0.5 | 0.4 | 0.3 | 0.2 | 0.1 | 0 |
|  | C | 0 | 0.2 | 0.4 | 0.6 | 0.8 | 1 | 0.8 | 0.6 | 0.4 | 0.2 | 0 |
|  | D | 0 | 0.1 | 0.2 | 0.3 | 0.4 | 0.5 | 0.6 | 0.7 | 0.8 | 0.9 | 1 |
| RNA survival rate | A | 0.5 | 0.5 | 0.5 | 0.5 | 0.5 | 0.5 | 0.5 | 0.5 | 0.5 | 0.5 | 0 |
|  | B | 1 | 0.9 | 0.8 | 0.7 | 0.6 | 0.5 | 0.4 | 0.3 | 0.2 | 0.1 | 0 |
|  | C | 0.1 | 0.2 | 0.4 | 0.6 | 0.8 | 1 | 0.8 | 0.6 | 0.4 | 0.2 | 0 |
| RNA level at age 0<sup>a</sup> | Unregulated transcription: 100 per time with no limit imposed by the DRA. |  |  |  |  |  |  |  |  |  |  |  |
|  | If TRA<DRA, then the RNA level at age 0=100; else RNA level at age 0=0. |  |  |  |  |  |  |  |  |  |  |  |
| DRA | Stable values: 50, 80, 150, 400 |  |  |  |  |  |  |  |  |  |  |  |
|  | Cycling change: alternating between 50 and 150 |  |  |  |  |  |  |  |  |  |  |  |
| | Aperiodic change: $DRA=400/(1+e^{4-0.2*time})$ | | | | | | | | | | | |

<sup>a</sup>TRA is total RNA activity at all RNA ages in the cell. DRA is the demand of RNA activity in the cell.

**Table S3.** Parameter values in the model to simulate the relationship between RNA and protein dynamics

|  | Age of protein or RNA<sup>a</sup> |  |  |  |  |  |  |  |  |  |  |
| --- | --- | --- | --- | --- | --- | --- | --- | --- | --- | --- | --- |
|  | 0 | 1 | 2 | 3 | 4 | 5 | 6 | 7 | 8 | 9 | 10 |
| RNA activity coefficient | 0 | 0.6 | 0.7 | 0.8 | 0.9 | 1 | 0.9 | 0.8 | 0.7 | 0.6 | 0 |
| RNA survival rate | 1 | 0.9 | 0.8 | 0.7 | 0.6 | 0.5 | 0.4 | 0.3 | 0.2 | 0.1 | 0 |
| RNA level at age 0<sup>b</sup> | If TRA < DRA, then RNA level at age 0=100; else RNA level at age 0=0. |  |  |  |  |  |  |  |  |  |  |
| DRA<sup>c</sup> | If TPA < DPA, DRA= (DPA-TPA)×2; else DRA=0. |  |  |  |  |  |  |  |  |  |  |
| Protein level/RNA activity<sup>d</sup> | 1 |  |  |  |  |  |  |  |  |  |  |
| Protein activity coefficient | 0 | 0.6 | 0.7 | 0.8 | 0.9 | 1 | 0.9 | 0.8 | 0.7 | 0.6 | 0 |
| Protein survival rate | 1 | 0.9 | 0.8 | 0.7 | 0.6 | 0.5 | 0.4 | 0.3 | 0.2 | 0.1 | 0 |
| Protein level at age 0 | If TPA < DPA and TRA ≥ 50, protein level at age 0=50; if TPA < DPA and TRA < 50, protein level at age 0=TRA; if TPA ≥ DPA, protein level at age 0=0. |  |  |  |  |  |  |  |  |  |  |
| DPA | To show the fluctuations of both RNA and protein levels: DPA=25 |  |  |  |  |  |  |  |  |  |  |
|  | To show the fluctuations of RNA levels and stable trends in protein levels: DPA=175 |  |  |  |  |  |  |  |  |  |  |
|  | To show the stable trends of both RNA and protein levels: DPA=200 |  |  |  |  |  |  |  |  |  |  |
|  | Cycling change: DPA alternating between 50 and 150 |  |  |  |  |  |  |  |  |  |  |

<sup>a</sup> Protein lifespan and age classification were set to be the same as those of RNA.
<sup>b</sup>TRA is total RNA activity at all RNA ages in the cell. DRA is the demand of RNA activity in the cell.
<sup>c</sup>TPA is total protein activity at all protein ages in the cell. DPA is the demand of protein activity in the cell. Here, the settings for the DRA depended on the values of the protein level/RNA activity ratios, protein survival rates, and protein activity coefficients. If TPA < DPA, protein age 1 was the first age to respond to the DPA shortage because the protein activity coefficient at protein age 0 was 0. The relationship between the DRA and protein activity at protein age 1 was: protein activity ÷ (protein level/RNA activity × protein survival rate at age 1 × protein activity coefficient at age 1) = protein activity ÷ (1 × 0.9 × 0.6) ≈ 2 RNA activity. Therefore, 2 was used in the equation, and it meant that 1 unit of insufficient protein activity could be transformed into 2 units of RNA activity.
<sup>d</sup>It meant that 1 unit of RNA activity could be transformed into 1 unit of protein level.

